# Nonmotor symptoms associated with progressive loss of dopaminergic neurons in a mouse model of Parkinson’s disease

**DOI:** 10.1101/2023.01.23.525182

**Authors:** Anna Radlicka, Judyta Jabłońska, Michał Lenarczyk, Łukasz Szumiec, Zofia Harda, Monika Bagińska, Joanna Pera, Grzegorz Kreiner, Daniel Wójcik, Jan Rodriguez Parkitna

## Abstract

Parkinson’s disease (PD) is characterized by three main motor symptoms: bradykinesia, rigidity and tremor. PD is also associated with diverse nonmotor symptoms that may develop in parallel or precede motor dysfunctions, ranging from autonomic system dysfunctions and impaired sensory perception to cognitive deficits and depression. Here, we examine the role of the progressive loss of dopaminergic transmission in behaviors related to the nonmotor symptoms of PD in a mouse model of the disease (the TIF-IA^DATCreERT2^ strain). We found that in the period from 5 to 12 weeks after the induction of a gradual loss of dopaminergic neurons, mild motor symptoms became detectable, including changes in the distance between paws while standing as well as the step cadence and sequence. Male mutant mice showed no apparent changes in olfactory acuity, no anhedonia-like behaviors, and normal learning in an instrumental task; however, a pronounced increase in the number of operant responses performed was noted. Similarly, female mice with progressive dopaminergic neuron degeneration showed normal learning in the probabilistic reversal learning task and no loss of sweet-taste preference, but again, a robustly higher number of choices were performed in the task. In both males and females, the higher number of instrumental responses did not affect the accuracy or the fraction of rewarded responses. Taken together, these data reveal discrete, dopamine-dependent nonmotor symptoms that emerge in the early stages of dopaminergic neuron degeneration.

## Introduction

The primary motor symptoms of Parkinson’s disease (PD) are caused by progressive degeneration of dopaminergic neuron projections from the ventral midbrain to the basal ganglia and the resulting imbalance in the activity of the direct and indirect striatal pathways^1,2^. PD is also associated with a variety of nonmotor symptoms that include impaired sensory perception^3–6^, autonomic dysfunction^7–10^, cognitive impairments, dementia and depression^11–14^. The wide diversity of the nonmotor symptoms is linked to the stages of development of PD, with neuron degeneration initially affecting the olfactory bulb in the brain and the enteric nervous system before advancing onto the sympathetic system, then spreading through the hindbrain, causing degeneration of noradrenergic and dopaminergic neurons, and eventually reaching the forebrain in the late stages^11,15,16^. Accordingly, the loss of olfactory acuity may precede motor symptoms and may even serve as an early indicator for diagnosis^17–19^. Moreover, in the later stages of the disease, the majority of PD patients develop dementia^13^, which is in line with this concept. This dementia is primarily related to a great load of Lewy bodies in the neocortical areas but could also be attributed to degeneration of cholinergic neurons within the nucleus basalis^20^.

Thus, PD is a multisystem disease^21^ that affects several neurotransmitter systems, including the dopaminergic, noradrenergic, serotonergic^22^ and cholinergic^23,24^ systems. The extent to which the loss of dopamine contributes to specific nonmotor symptoms is varied and in many cases remains controversial. For instance, while cognitive deficits in the later stages of PD were primarily attributed to cholinergic degeneration and thus resemble the symptoms of dementia associated with Alzheimer’s disease, an impairment of executive functions is reportedly specific to PD and includes deficits in attention, impaired problem solving and action sequencing^25^. These symptoms are also reported in PD without dementia^26^ and could be plausibly attributed to impaired dopamine signaling in the basal ganglia. Executive dysfunctions, already present in a considerable fraction of newly diagnosed patients^27^, may stem from dopamine deficits that affect fronto-striatal loops, while PD dementia that emerges in later PD stages could be caused by the impairment of other neurotransmitter signaling^28^. Although neuroimaging evidence points to the dopaminergic origins of executive dysfunctions in PD^29^, the correlation of their severity with cardinal motor symptoms, also dependent on dopamine signaling, gives mixed results between the studies^30–32^. Moreover, the incidence of depression is significantly increased in the prodromal phase of PD and correlated with altered hyperechogenicity of the substantia nigra^33^, which again points to altered dopaminergic transmission as a potential cause. However, in the case of either executive impairments or depression, the degeneration of the locus coeruleus noradrenergic neurons could be argued to be an equally plausible cause, as could the impact of disrupted cholinergic transmission. Finally, limited correlations exist in the development of motor and nonmotor symptoms^34–36^, and attempts to classify PD subtypes on the basis of the cooccurrence of both types of symptoms remain inconclusive^37^.

Therefore, to isolate the contribution of dopaminergic neurons in the development of nonmotor symptoms of PD, the model of choice is genetically modified mice with an inducible loss of dopaminergic neurons by the use of the Cre-loxP system that targets functionally essential genes in DAT-expressing neurons, such as in MitoPark^38^ and TIF-IA^DATCreERT2^ strains^39^. The use of these mice allows us to study the Parkinsonian phenotype that develops relatively slowly (compared to neurotoxic models) and is selectively caused by dopaminergic dysfunction. Although these models have, arguably, limited construct validity (i.e., the underlying cause of the loss of neurons is not the same as in PD), they were shown to accurately recapitulate motor impairment^38–41^, selected nonmotor symptoms of PD^42^ and exhibit at least some cell loss mechanisms similar to that of PD, i.e., mitochondrial dysfunction or nucleolar disruption. Therefore, they proved valuable in investigating the consequences of the loss of dopaminergic neurons and testing new potential treatments for PD. Here, we use the TIF-IA^DATCreERT2^ strain to assess the development of motor and nonmotor symptoms in the early and middle stages of dopaminergic neuron degeneration. We have tested in parallel the emergence of changes in gait together with symptoms related to impaired olfaction, anhedonia-like behavior and cognitive performance.

## Materials and methods

### Animals

Mice were housed at the animal facility of the Maj Institute of Pharmacology of the Polish Academy of Sciences under a 12/12 hour light/dark cycle at 22±2 °C and 55±10% air humidity. Males were housed in groups of 2-4 siblings per cage, except for saccharin preference test sessions, during which they were housed individually for periods of 24 hours. Females were initially housed 2-4 per cage and were later moved to two IntelliCages, 11 females in each. Animals had *ad libitum* access to food and water, except for male mice assessed for olfactory acuity (buried food task), which were deprived of food access for 24 h prior to the test.

Male and female TIF-IA^DATCreERT2^ [Tg/0; flox/flox] mice were generated as described previously^39,40^. Control animals lacked the CreERT2 transgene ([0/0; flox/flox] genotype). *Tif1a* inactivation in dopamine transporter (DAT)-expressing cells was induced in 9- to 11-week-old male and female mice by tamoxifen treatment (Sigma, Germany); specifically, 2 mg dissolved in 100 μl of sunflower oil was injected once daily *i*.*p*. for 5 consecutive days. Control mice also received tamoxifen treatment. All mice were allowed to rest for 3 weeks before testing started. Experimental procedures were performed on 22 males (10 mutants and 12 controls) and 22 females (9 mutants and 13 controls), including 1 control female that was excluded from the probabilistic reversal learning test in the early adaptation phase.

All procedures were conducted in accordance with the ARRIVE guidelines, Polish law and European Commission regulations concerning the care and use of laboratory animals (Directive 2010/63/UE, European Convention for the Protection of Vertebrate Animals Used for Experimental and other Scientific Purposes ETS No.123, and Polish Law Dz.U. 2015 poz. 266). All experimental procedures were approved by the II Local Institutional Animal Care and Use Committee in Krakow (permits no. 28/2019, 197/2019 and 198/2019).

### Behavioral procedures

Male and female mice underwent regular testing for gait parameters and behaviors related to nonmotor Parkinson’s disease symptoms starting on the 4^th^ week after tamoxifen injection. A schematic representation of the experiment is shown in Figure 1. In addition to gait analysis, male mice were tested for operant sensation seeking, olfactory acuity and saccharin preference. Female mice remained in an IntelliCage apparatus throughout the procedure (except for the gait recording sessions) and were tested for probabilistic reversal learning. The body weights of all animals were also monitored.

**Figure 1.**
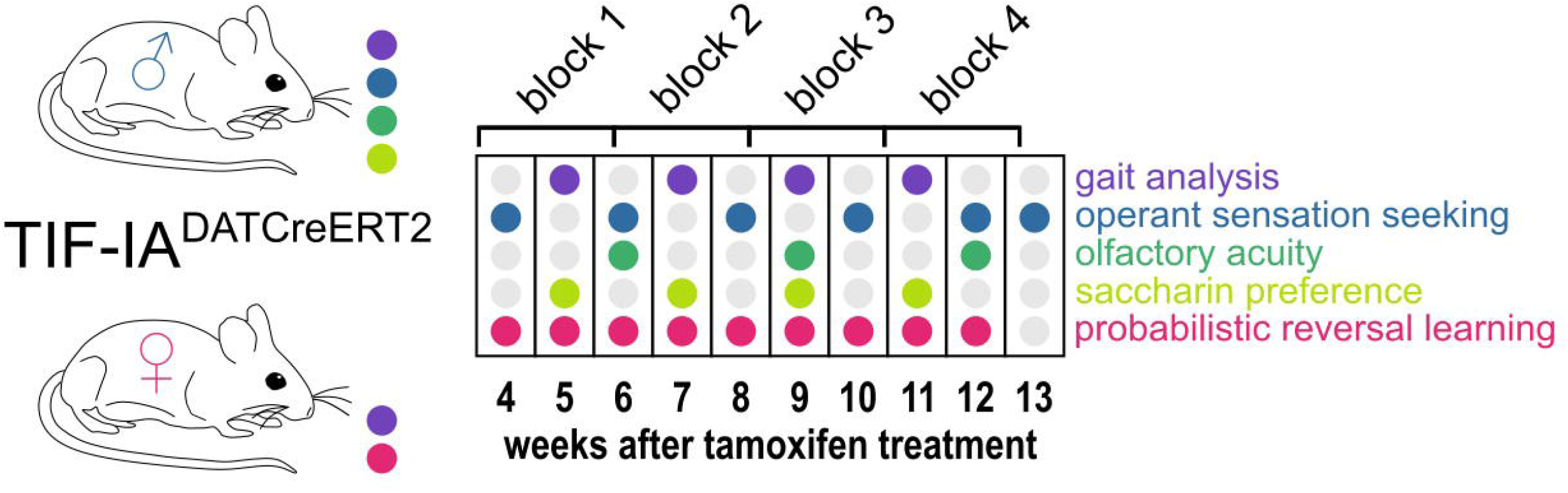
Experimental design. The diagram shows the schedule of gait (Catwalk, violet dots) and nonmotor behaviors tested (all remaining colors, as indicated on the left) in male and female TIF-IA^DATCreERT2^ mice and controls.

### Gait

All mice were tested for gait on the 5^th^, 7^th^, 9^th^ and 11^th^ week after tamoxifen injection using the CatWalk XT system v10.6 (Noldus, the Netherlands). The tests were performed during the light phase. Briefly, mice were placed individually at either of the two ends of a tunnel covering a side-lit glass platform and were allowed to move freely in the tunnel. A camera placed under the platform recorded the body movement and paw placement, and the recordings were validated by the software. A total of 2-6 runs of each mouse per test session were used for statistical analyses. Recordings that did not include at least 1 complete pattern detected were excluded from the analysis.

### Operant sensation seeking

The operant sensation seeking test was performed as described previously^43–45^. Male mice were placed individually in operant conditioning chambers ENV-307W (Med Associates, USA). During the 1-hour sessions, the animals were able to freely explore the cage and perform active and inactive operant responses by placing their snouts in one of the two holes located ∼2 cm above the grid floor. The active operant response resulted in the presentation of a blinking light with a randomized duration of 2, 4, 6 or 8 s, a frequency of 0.625, 1.25, 2.5, or 5 Hz and a 2.9 kHz 65 dB monotone beep (ENV-323AW Sonalert, Med Associates). The assignment of active and inactive operant holes was fixed for each mouse throughout the experiment. The number of active and inactive operant responses was recorded by the software.

### Olfactory acuity

Olfactory acuity was assessed in the buried food test as described by Yang and Crawley^46^. One to three days before each test, mice received vegetable-flavored crackers in their home cages (“Sunbites” brand, Frito Lay Poland). All food was removed from the home cage on the day prior to the test. Olfactory acuity was assessed by placing the mice individually in an opaque cage (interior dimensions: 53 × 32 × 19 cm) filled with a 3-cm layer of aspen bedding. First, a mouse was able to explore the cage freely, and after acclimatization, it was briefly transferred to a separate cage while a cracker (approx. 2 g) was buried beneath the bedding in one of the corners. The mouse was then reintroduced to the opposite corner of the cage. The observer recorded the delay before the mouse started digging in the correct corner and the time to retrieve the cracker (i.e., digging out or taking a bite). The location of the cracker was randomized and changed after each completed trial.

### Saccharin preference

Saccharin preference was measured as described previously^47^ with minor modifications. Males were placed in individual small cages (26 × 20 × 135 cm) for 24 hours and were able to drink tap water or a 0.1% (w/v) saccharin solution (Sigma Aldrich, Germany) from measuring cylinders. Fluid consumption was assessed based on the change in liquid volume. The position of the cylinders was switched between consecutive tests. Changes in volume ≥ 20 ml were assumed to result from leaks and were excluded from analysis.

### Probabilistic reversal learning

The probabilistic reversal learning task was performed as described previously^48^. Briefly, female TIF-IA^DATCreERT2^ mice were implanted subcutaneously with PICO UNO radio-frequency identification (RFID) transponders 4 weeks after mutation induction and placed in groups of 11 in two automated IntelliCage systems (TSE Systems, Germany). Each group included both genotypes. The cages recorded all entries of an animal into a corner and counted the number of licks on each of the bottles placed in the corners. Each corner was fitted with two 250-ml bottles, accessible through ports with access controlled by sliding doors. Food was available *ad libitum* throughout the procedure. At the start of the procedure, mice were introduced into the IntelliCage and allowed free access to all the drinking bottles (i.e., the sliding doors were in the open position). Next, bottles in two of the corners were replaced with 0.1% (w/v) saccharin solution, and a restriction was placed on access: the doors blocking the access to bottles opened for 10 s after the animal was detected in the corner, with a 0.5 s delay in the case of water bottles and a 2 s delay for saccharin bottles. The bottles filled with saccharin solution could be accessed by mice in 90% of visits, while water remained fully accessible. The position of the saccharin and water bottles was changed every 48 h to avoid corner preference. After 3 position changes, the probabilistic reversal phase started. The position of the corners with saccharin bottles was fixed, but access to the bottles was granted with 90% probability in one of the corners and 30% in the other. The probabilities were reversed every 48 h, and the experiment continued until the end of 12 weeks after the mutation was induced. The procedure was interrupted several times, e.g., for brief periods of several hours required for gait testing, weight measurements, cage cleaning and maintenance. Periods of time with interruption of correct system operation were excluded from the analysis (a complete list is provided in the script used for the statistical analysis).

### Immunofluorescence staining

On the 123rd day (18^th^ week) after tamoxifen treatment, the animals were killed by cervical dislocation. Their brains were extracted and fixed in 4% paraformaldehyde (Roth, Germany) solution at 4°C for two days, and the fixative was then changed to 0.4% paraformaldehyde. After 55 days, the brains were stored in phosphate-buffered saline solution (PBS; Serva, Germany) with 0.05% thimerosal (Sigma). The brains were sectioned on a Leica VT1200 (Leica, Germany) vibratome into 40-µm coronal sections, including the olfactory bulbs (bregma 4.28 to 3.92 mm according to the mouse brain atlas by Paxinos and Franklin^49^), forebrain (bregma 1.54 to 0.14 mm) and midbrain (bregma -2.92 to -3.16 mm). The sections were collected in 48-well plates filled with 150 µl of PBS and refrigerated until staining.

The staining procedure was performed in accordance with a previously published protocol^50^. After washing in PBS with 0.2% Triton X-100 (Amresco, USA), the sections were blocked for 30 minutes in 5% normal pig serum (Vector Laboratories, USA) in PBS with 0.2%-Triton X-100 and then incubated overnight at 4 °C with primary antibodies (1:500; Anti-Tyrosine Hydroxylase Antibody Sigma⍰Aldrich, USA, catalog no. AB1542) in the same blocking solution. The sections were then rinsed with PBS-Triton X-100 solution and incubated for 1 hour with the secondary antibody (1:100; rabbit anti-sheep IgG antibody (H+L), fluorescein, Vector Laboratories, cat. no. FI-6000), rinsed again with PBS and mounted with VECTASHIELD^®^ HardSet™ Antifade Mounting Medium with DAPI (Vector Laboratories, cat. no. H-1500). Images were captured with a Leica TCS SP8 X confocal laser-scanning microscope (Leica Microsystems, Germany) using HC PL FLUOTAR 10x (0.30) and 40x (0.60) objectives and Leica Application Suite X software (Leica Microsystems).

### Statistical analysis

CatWalk data were first reduced to exclude variables without variance, missing measurements, and variables that were directly influenced by animals’ speed, since they may have been influenced by experimenters. Next, the corresponding parameters collected from the left and right paws were correlated and averaged, and parameters that had weak correlation (Pearson correlation coefficient R < 0.5) were excluded from further analyses, as they introduced noise into the analysis. To extract gait measures of interest that were potentially affected by partial loss of dopamine neurons, a logistic regression model, glm (formula = genotype ∼ sex + days + parameter, family = binomial), was then applied. A stepwise variable selection algorithm was used to select variables relevant for the prediction of genotype from data, starting from a null model comprising only date and sex as predictors.

The parameters included in the final regression model as a result of the selection algorithm were further analyzed in R v4.0.4. Both CatWalk data and measurements collected from tests evaluating nonmotor functions and body weight were individually subjected to a mixed two-way analysis of variance (ANOVA) to determine the effects of mutation and the time elapsed and their interaction. Animals that lacked data in at least one time point for a variable were excluded from the analysis of that particular parameter. Additionally, Bonferroni-adjusted t-tests were carried out to assess differences between the groups at each time point as *post hoc* analyses. The analyses for males and females were carried out separately.

The correlation analysis of motor and nonmotor function measurements was performed as follows: starting from the 23rd day after the last day of tamoxifen treatment, time points were grouped into four 16-day-long blocks. Blocks 2, 3 and 4, i.e., the results collected from the 39^th^ day after tamoxifen treatment onward, were considered in the analysis. Olfactory acuity results and female weight were excluded from the correlation analysis due to missing data. A |R| value of ≥ 0.4 was significant and corresponded to a *P* value of < 0.05 (Bonferroni adjusted for multiple parallel correlations).

## Results

### Loss of dopaminergic neurons in TIF-IA^DATCreERT2^ mice

The TIF-IA^DATCreERT2^ mouse strain is characterized by a progressive loss of dopaminergic (i.e., dopamine-transporter (DAT) expressing) neurons after the induction of the mutation with tamoxifen^39,40^. To confirm that the mutation caused a decrease in the midbrain DAT-positive cells consistent with that previously reported, the loss of dopaminergic neurons was assessed by immunofluorescence staining for tyrosine hydroxylase (TH, Fig. 2) in brains extracted 18 weeks after tamoxifen treatment (i.e., after all the behavioral tests had been completed). As shown in Fig. 2a, robust TH staining was observed in the dorsal striatum, the nucleus accumbens and the olfactory tubercle of control animals, i.e., structures with the most extensive dopaminergic innervation^51^, while the fluorescence signal was notably weaker in the adjacent cortical areas. Conversely, immunofluorescence in the corresponding striatal areas in a representative section from a TIF-IA^DATCreERT2^ mouse was weaker or absent and comparable to the signal observed in the dorsal cortical areas (Fig. 2b). Accordingly, a difference in TH staining was also observed at the level of the midbrain, where the number of TH-positive cell bodies was appreciably larger in control animals (representative example in Fig. 2c) than in mutants (Fig. 2d). In line with a previous report^52^ The DATCreERT2-driven mutation had no appreciable effects in the main olfactory bulb, where a similar intensity of fluorescence was observed in control (Fig. 2e) and mutant (Fig. 2f) animals. In both cases, intense TH immunofluorescence was located within the external, glomerular layer, which contains the dopamine- and GABA-releasing interneurons of area A16^53,54^. As shown in the close-up Figures 2g and 2h, the TH signal was present both in cell bodies and neurites in the glomerular layer, again with no appreciable differences between mutant and control animals. Thus, at 18 weeks after tamoxifen administration, the mutation caused an extensive loss of TH-positive cells in the ventral midbrain but had no observable effect on the abundance of putative dopaminergic cells in the olfactory bulb. The extent of the mutation within the striatal and midbrain areas was consistent with the originally reported analysis^39^.

**Figure 2.**
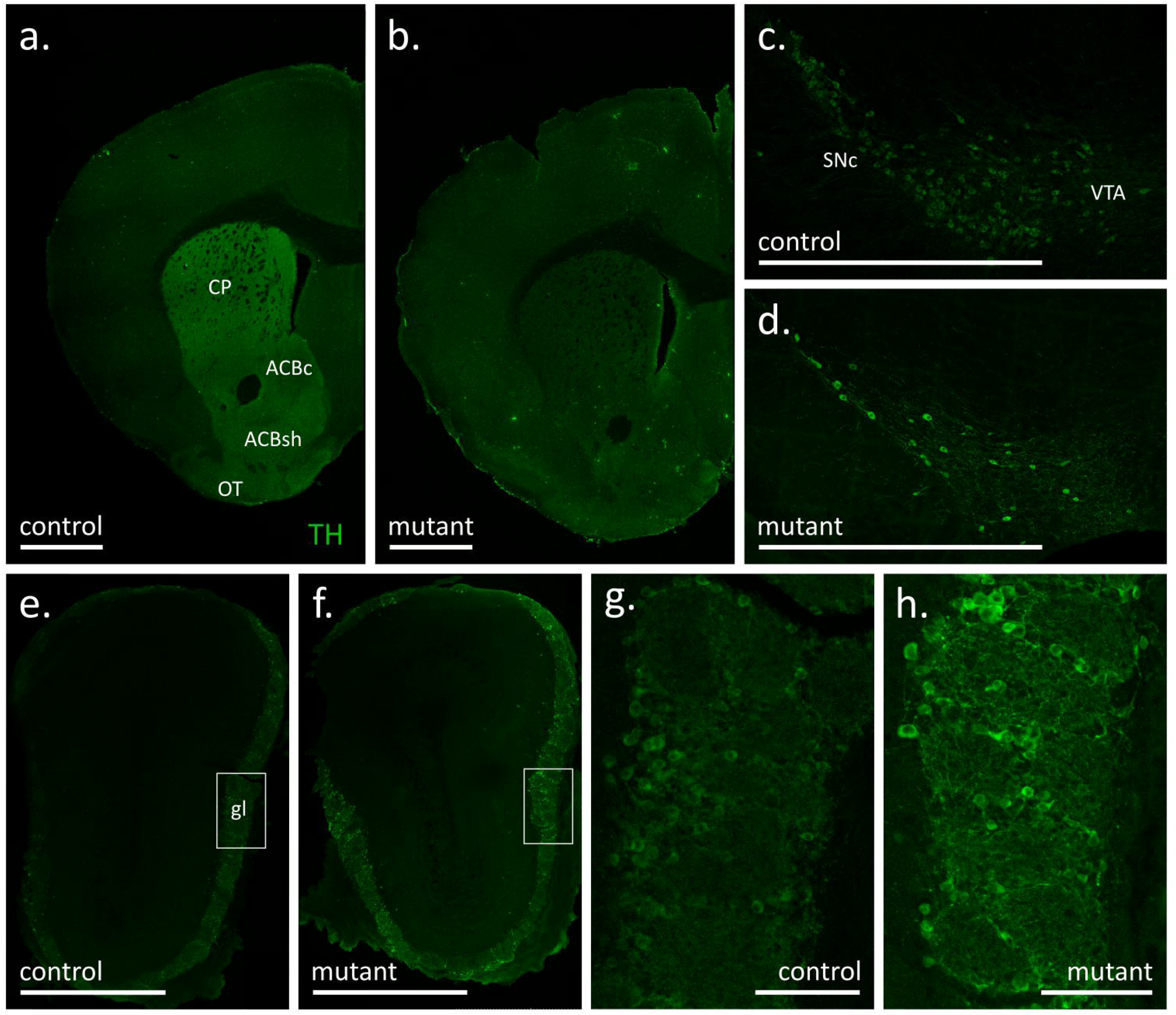
Tyrosine hydroxylase (TH) immunofluorescence signal in coronal sections of the forebrain (a-b), substantia nigra with ventral tegmental area (c-d) and olfactory bulbs (e-h) of control (a, c, e, g) and mutant (b, d, f, h) mice at 18 weeks after tamoxifen treatment. White bars correspond to 1 mm (a-f) or 100 µm (g-h).

Dopamine signaling is necessary for eating-oriented behavior. In humans, changes in weight were reported in PD patients in the premotor phase of the disease^55^. In mice, a complete loss of dopamine production is fatal, as animals do not consume sufficient amounts of food for sustenance^56^. Here, we found that TIF-IA^DATCreERT2^ males were on average 5% lighter than the controls and that this difference was observable even before the induction of the mutation (F_(1, 18)_ = 7.026, *P* = 0.0163; Fig. 3a; a full list of all ANOVAs for the performed tests and measurements is included in Supplementary Table S1). The difference in weight increased after the mutation was induced, as evidenced by the significant interaction of the genotype and time elapsed from tamoxifen administration (F_(5, 90)_ = 2.56, *P* = 0.0326). However, we note that despite the genotype, the weight continued to significantly increase over time in all animals (F_(5, 90)_ = 87.08, P < 0.0001). In females (Fig. 3b), body weight also increased over time (F_(3, 57)_ = 128.663, P < 0.0001), but the effects of the mutation (F_(1, 19)_ = 3.32, P = 0.0842) and interaction (F_(3, 57)_ = 1.773, P = 0.163) were not significant. Therefore, although all mice gained weight throughout the experiment, dopamine cell loss mildly affected the body weight of male mutants.

**Figure 3.**
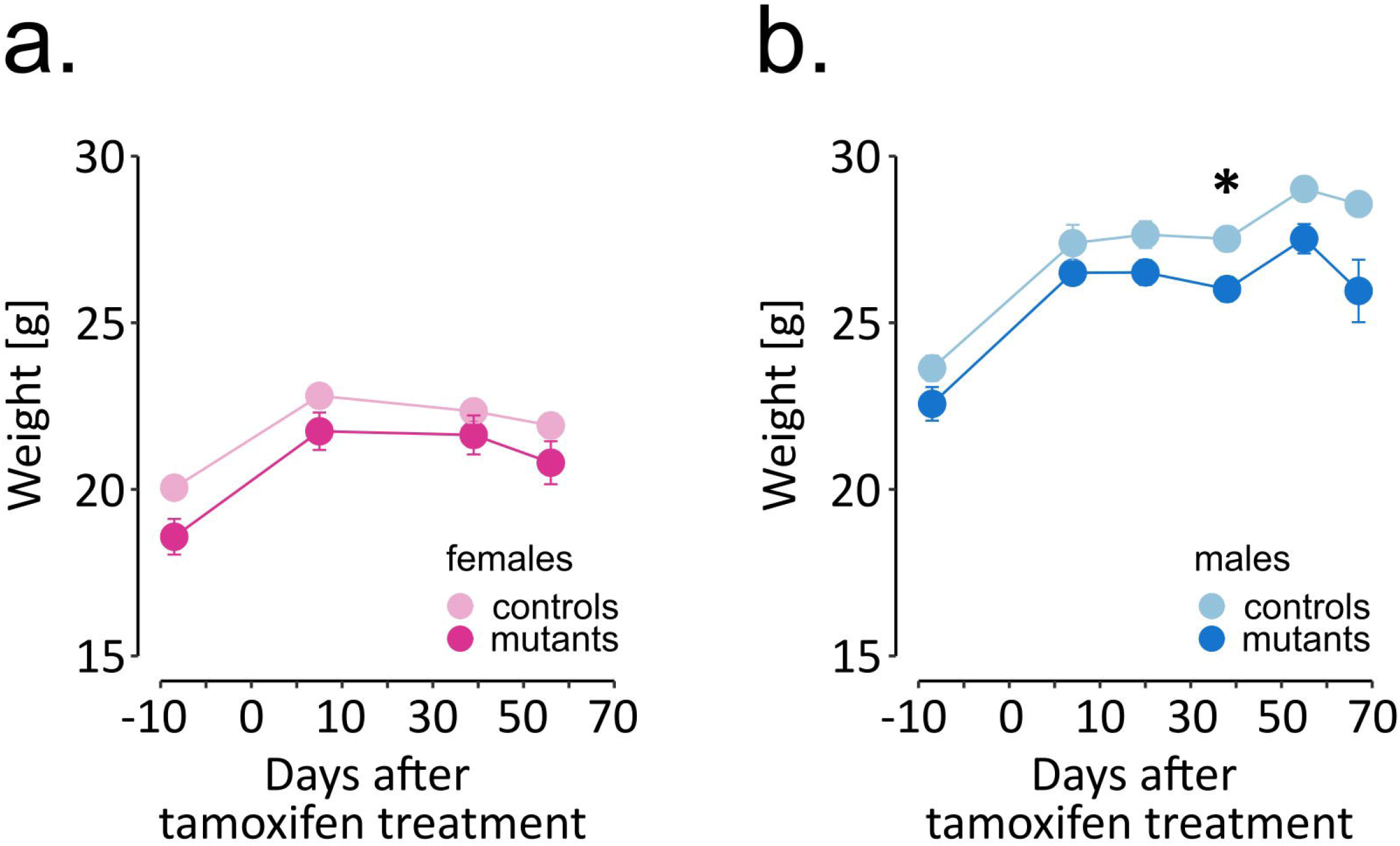
Weight of TIF-IA^DATCreERT^2 and control mice. (a, b) The graphs show the weight of the experimental animals immediately before tamoxifen treatment (−7 days time point) and throughout the testing procedure. Each point represents the group mean, and error bars are SEM. Female mice are shown in shades of pink, and male mice are shown in blue. Darker hues are TIF-IA^DATCreERT2^ mice, and lighter hues represent the controls. Stars indicate significant difference mutant vs. control, *t*-tests with Bonferroni correction for multiple tests, p <0.05.

### Effects of the mutation on motor function

To assess the effect of the developing loss of dopaminergic neurons on motor function in TIF-IA^DATCreERT2^ mice, we conducted gait analysis using the CatWalk system (a description of relevant gait characteristics and parameters is provided in Table 1). Animals were tested 4 times in 2-week intervals, starting on the 5^th^ week after the induction of the mutation. Over 200 raw parameters and derived measures were generated from each session (see Supplementary Table S2 for the complete raw dataset). Two-step data reduction was performed. First, we removed measures that were not recorded or were confounded by the way the procedure was performed. A total of 31 parameters with only zeroes or no values scored were discarded. Additionally, we removed 7 parameters directly related to animals’ speed because the experimenter had to intervene at the start of a considerable fraction of trials. Second, correlations between the left and right paws for the corresponding measures were assessed separately for the front and hind limbs (see Supplementary Table S3 for summary). All measures with Pearson R < 0.5 were excluded as having excessively high variance, and the remaining parameters were averaged between the paws. A total of 62 parameters remained after data reduction (Supplementary Table S4) and were used to construct a linear regression model with the fewest parameters that achieved the highest accuracy in predicting the genotype. The optimal model included 15 gait parameters and 3 days x parameter interactions (AIC = 700.3788, cross-validated accuracy = 0.688, Supplementary Table S5), primarily related to the hind paw print size, paw placement coordination and timing, and cadence. Thus, these 15 gait parameters were indicated to be most closely related to the animals’ genotype.

**Table 1.** Description of selected gait characteristics and parameters measured/analyzed by the CatWalk system in this study. Abbreviations: F – front paw, H – hind, L – left, R – right.

| Gait characteristic | Description | Example of CatWalk parameters [units] |
| --- | --- | --- |
| Stand or Stance | A phase of gait during which a paw is in contact with glass walkway/walking surface. | Stand [s] |
| Swing | A phase of gait during which a paw is not touching the walkway. | Swing speed [s] |
| Step cycle | Time between two consecutive initial contacts of the same paw and the walkway; it equals the sum of stand and swing times during a single step. | Step cycle [s] |
| Base of support | The average width between the two front or the two hind paws. | Base of support Front Paws [cm] |
| Cadence | Number of steps taken per second in a run; it equals:<br>$\frac{\text{number of steps} - 1}{\text{initial contact last Stand} - \text{initial contact first Stand}}$ | Cadence [number per second] |
| Couplings | A temporal relationship between placing two paws on the | Couplings RF -> LF [%] |
|  | walkway during one step cycle, i.e., at which moment is the second paw placed in relation to the time of placement of the first one. |  |
| Step sequence | An order of paw placement (footfall pattern). There are 6 footfall patterns (<u>A</u>lternate, <u>C</u>ruciate, <u>R</u>otate):<br>AA: RF -> RH -> LF -> LH<br>AB: LF -> RH -> RF -> LH<br>CA: RF -> LF -> RH -> LH<br>CB: LF -> RF -> LH -> RH<br>RA: RF -> LF -> LH -> RH<br>RB: LF -> RF -> RH -> LH | Step sequence AA<br>[% of total number of all footfall patterns in a run] |
| Max contact | A moment in which a paw makes a maximum contact with the glass walkway since the start of a run. | Max contact mean intensity<br>[value of pixels in the range from 0 to 255] |
| Print positions | A distance between the ipsilateral paws, where the front paw has been placed earlier than the hind paw in the same step cycle. | Print positions Right Paws [cm] |

The TIF-IA^DATCreERT2^ mice differed from controls in several parameters that describe the gait characteristics of hind paws (Fig. 4; a full list of ANOVA results is included in Supplementary Table S1). The progressive loss of dopamine neurons affected the gait of males and females to varying degrees. The base of support of the hind paws (two-way ANOVA; males, genotype: F_(1, 17)_ = 4.972, *P* = 0.0395; Fig. 4a), their print area (F_(1, 17)_ = 5.276, *P =* 0.0346; Fig. 4c), print width (F_(1, 17)_ = 4.485, *P =* 0.0492; Fig. 4e) and the interval (‘couplings’) between the placement of the right hind (RH) and the left front (LF) paws within a step cycle (F_(1, 16)_ = 16.73, *P =* 0.000855; Fig. 4g) were significantly altered in mutant males but not mutant females (Fig. 4b, d, f and j, respectively). These 4 parameters were, on average, smaller in value in mutant males than in their control littermates. Conversely, hind paw swing speed was not significantly affected by the mutation in males (Fig. 4i), but its effects were significant in females (F_(1, 17)_ = 5.535, *P =* 0.0309; Fig. 4j), with TIF-IA^DATCreERT2^ females having higher swing speed than controls. Significant interactions between time and genotype, the hind paw print width in females only (females: Fig. 4f, F_(3, 51) =_ 2.826, *P* = 0.0478; Fig. 4e for males) and couplings RH - > LF were only observed in two cases in males (Fig. 4g, F_(3, 48)_ = 3.161, *P* = 0.0329) but not females (Fig. 4h).

**Figure 4.**
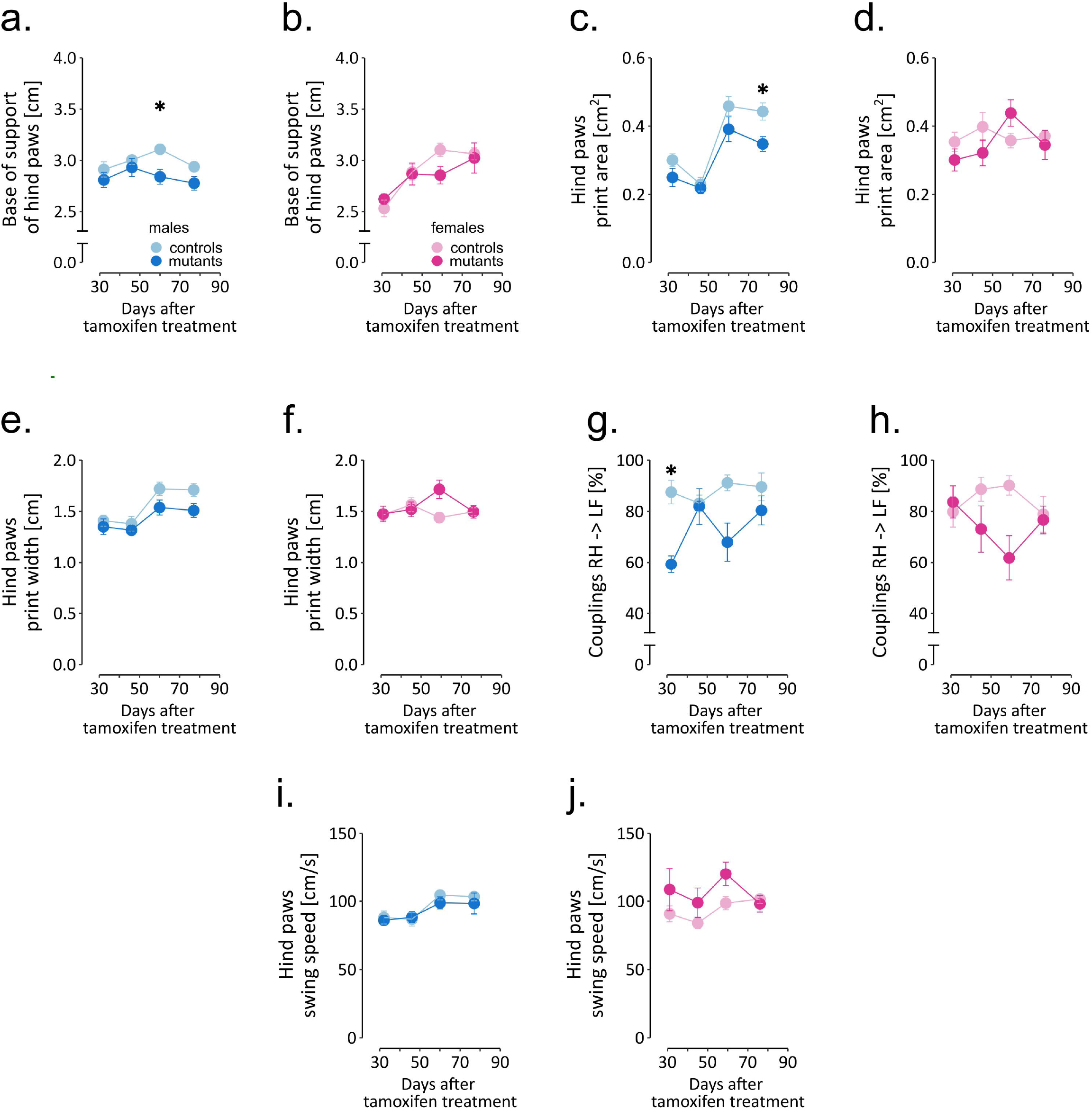
Selected gait parameters in TIF-IA^DATCreERT2^ and control mice. a to e summarize measurements of the five selected parameters in female mice, and f to j correspond to male mice. Each point represents the group mean, and error bars are SEM. Female mice are shown in shades of pink, and male mice are shown in blue. Darker hues are TIF-IA^DATCreERT2^ mice, and lighter hues represent the controls. Stars indicate significant difference mutant vs. control, *t*-tests with Bonferroni correction for multiple tests, test p <0.05. Abbreviations: BOS – base of stand, AA – alternate footfall pattern with the paw sequence: RF -> RH -> LF -> LH, RF – right front paw, LF – left front.

Additionally, the effects of time were significant in several of the parameters (Supplementary Table S1) that describe hind paw print size by print area (males: days: F_(3, 51)_ = 44.889, *P* < 0.0001; Fig. 4c for males and d for females) and max contact mean intensity of pixels (males: F_(3, 51)_ = 12.719, *P* < 0.0001; Supplementary Fig. S1a for males and b for females), paw placement coordination in couplings RF -> RH (females: F_(3, 51)_ = 4.425, *P* = 0.0077; Supplementary Fig. S1c and d) and alternate step sequence footfall pattern AA (males: F_(3, 51)_ = 3.068, *P =* 0.036; females: F_(3, 51)_ = 13.768, *P* < 0.0001; Supplementary Fig. S1e and f), swing speed (males: F_(3, 45)_ = 7.481, *P =* 0.0004; Fig. 4i and j), step cycle (males: F_(3, 45)_ = 7.240, *P =* 0.0005; females: F_(3, 51)_ = 7.241, *P =* 0.0004; Supplementary Fig. S1g and h) and cadence (males: F_(3, 51)_ = 7.054, *P =* 0.0005; females: F_(3, 51)_ = 4.088, *P =* 0.0112; Supplementary Fig. S1i and j) and base of support of the front paws (males: F_(3, 51)_ = 11.428, *P* < 0.0001; Supplementary Fig. S1k and l). Without significant interaction, these changes likely reflect age effects on movement related to weight gain. Of these parameters, cadence, alternate step sequence footfall pattern AA, hind paw max contact mean intensity of pixels in males and hind paw base of support in females displayed a tendency to increase over time, while hind paw step cycle tended to decrease and the others fluctuated or remained relatively stable between the consecutive test sessions. ANOVA did not detect significant effects in the remaining parameters implied by the regression model (Step Sequence CB, Step Sequence RB, Couplings RF -> LF and Print Positions Right Paws; Supplementary Table S5).

Taken together, these results show that in the period between 5 and 12 weeks after the mutation was induced, the effects on gait remained at most moderate, and only relatively minor impairments were observed in TIF-IA^DATCreERT2^ mice. The most consistent effects of dopaminergic neuron loss were observed in the parameters related to body weight distribution on the hind paws in male mutants.

### Impact of dopaminergic cell loss on nonmotor behaviors in males

In parallel to the gait analysis, we assessed nonmotor behaviors potentially relevant to known early PD symptoms. Male TIF-IA^DATCreERT2^ mice were tested at 2- to 3-week intervals for olfactory acuity, saccharin preference and operant sensation seeking (OSS; complete data are presented in Supplementary Table S6). In the OSS task, mice were offered two operands (nose pokes), and the response to one of them was associated with the presentation of a semirandom sequence of blinking lights and tones. C57Bl/6 mice were observed to readily learn to perform responses associated with sensory stimuli, which is interpreted as a sensation-seeking-like behavior^43,44^. Importantly, the task did not require food or drink deprivation, which would be a major confounding factor in the case of a model animal with a progressive loss of dopaminergic neurons. We observed that while all mice showed an increase in instrumental responses on the operant associated with the semirandom sensory stimulus over time (Fig. 5a, F_(16, 288)_ = 17.155, *P* < 0.0001), a significant interaction was also observed between time and genotype (F_(16, 288)_ = 2.844, *P* = 0.0003), but genotype did not have a base effect (F_(1, 18)_ = 1.452, *P* = 0.244). On average, the mutants performed 73.9 ± 5.93 active nose pokes per session, while controls only performed 49.6 ± 4.04 nose pokes. Conversely, the number of responses on the “inactive” operant remained low, without genotype (F_(1, 18)_ = 0.53, P = 0.476) or interaction (F_(16, 288)_ = 1.378, P = 0.151) effects, although a significant effect of time was observed (Fig. 5b, F_(16, 288)_ = 2.137, *P* = 0.007).

**Figure 5.**
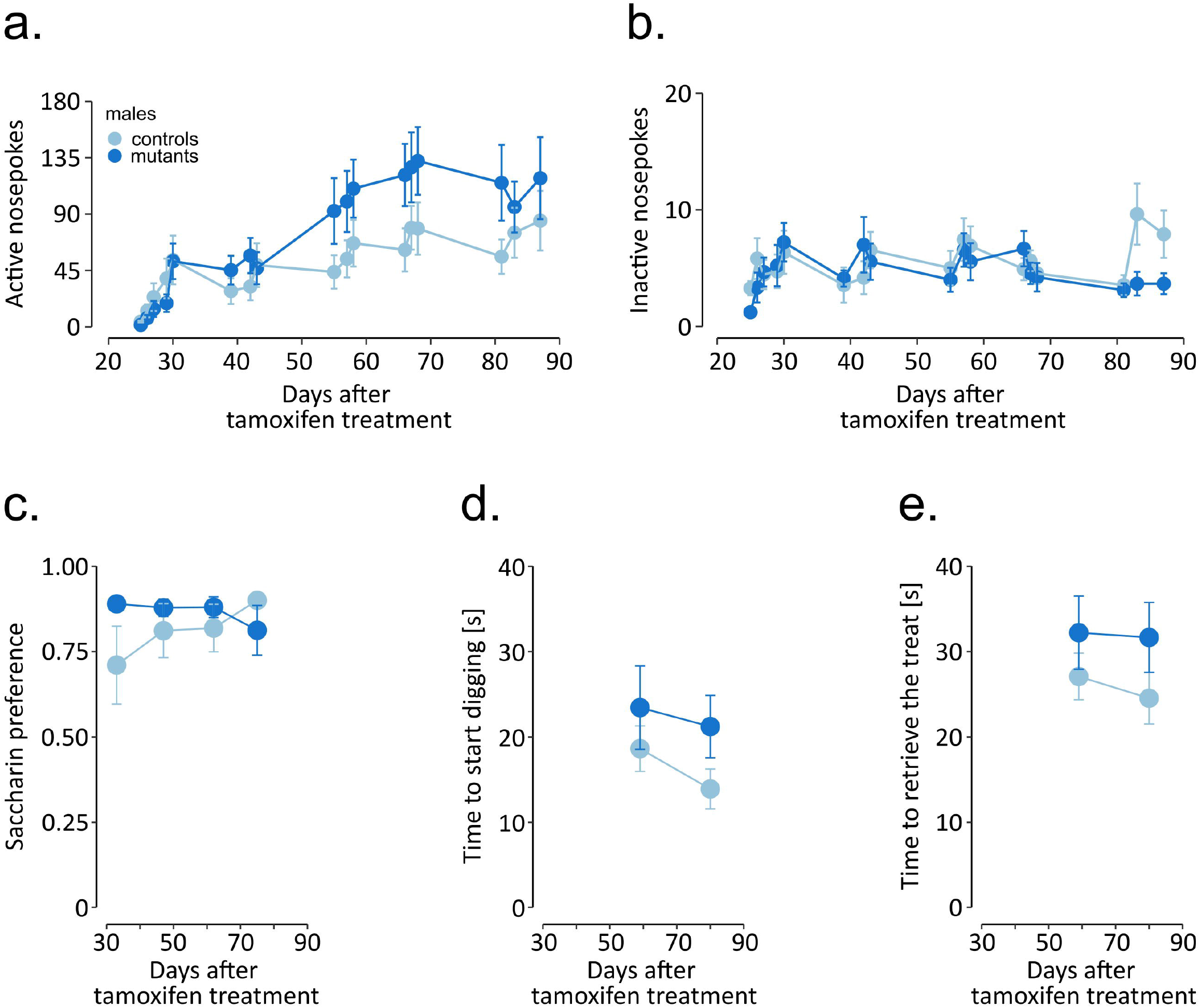
Nonmotor behaviors in male mice. The graphs show (a, b) instrumental responding in the OSS task, (c) two-bottle choice saccharin preference, and (d, e) time to find and retrieve a treat in the olfactory acuity test. Points represent the group mean, and error bars are SEM. TIF-IA^DATCreERT2^ mice are shown in dark blue, and the lighter color represents the controls. Stars indicate significant difference mutant vs. control, *t*-tests with Bonferroni correction for multiple tests, p <0.05.

To test whether the male mutants also displayed a higher (pleasant sensation seeking) or lower (anhedonia-like behavior) propensity for another kind of reward, i.e., sweet drinking liquid, they were tested for saccharin preference every two weeks, starting 5 weeks after tamoxifen administration. Both mutants and controls exhibited similarly high preference toward drinking saccharin in comparison to water (Fig. 5c), and the mutation had no significant effect on the observed preference (F_(1, 15)_ = 1.29, *P* = 0.274, two-way ANOVA).

Next, we tested whether dopaminergic cell loss affected their sense of smell. The olfactory acuity test was performed once every three weeks, starting 6 weeks after tamoxifen administration; the first session was treated as training and excluded from the analysis. Each time, the cracker was hidden in a different location, and mutants and controls exhibited a similar time to start digging at the ‘correct’ corner, i.e., where the cracker was buried underneath the bedding (F_(1, 18)_ = 2.418, *P* = 0.137, two-way ANOVA; Fig. 5d). Additionally, we also recorded the time in which the mice retrieved the cracker, and mutation did not significantly affect this time (F_(1, 18)_ = 3.121, *P* = 0.094; Fig. 5e).

Dopaminergic cell loss in TIF-IA^DATCreERT2^ males led to increased active operant responses over time as a measurement of sensation seeking. At the same time, the male mutants displayed lower counts of inactive responses, similar to those exhibited by the controls. These observations show that the mutants did not present learning or memory deficits and that their behavior was mainly directed toward receiving stimuli.

### *Probabilistic reversal learning in female* TIF-IA^DATCreERT2^ mice

Female mice were tested for executive function impairments in a probabilistic reversal learning task based on the IntelliCage system. In this version of the task, probabilistic choices are performed in the home cage, without any form of deprivation, and in group-housed animals. Animals were first habituated to the cage and completed an adaptation phase. The probabilistic choice task started on the 5^th^ week after the mutation was induced. Data were binned in 16-day periods centered around the three latter time points from the gait analysis (Fig. 6; data in 24 h bins are summarized in Supplementary Table S7). Female TIF-IA^DATCreERT2^ mice made more visits per day in the drinking compartments than controls (F_(1, 19)_ = 10.86, *P* = 0.0044; Fig. 6a), especially in the first two periods. Additionally, a significant effect of time (F_(2, 38)_ = 13.129, *P* < 0.001) and a significant interaction of time and genotype (F_(2, 38)_ = 3.954, *P =* 0.028) were observed. The number of choices (visits longer than 2 s in the compartments offering only saccharin) followed the same course, with a significant increase in mutants across all time periods (F_(1, 19)_ = 16.34, *P* < 0.001; Fig. 6b) and a genotype-independent effect of time (F_(2, 38)_ = 6.252, *P* = 0.005), but no significant interaction (F_(2, 38) =_ 0.281, *P =* 0.757). Genotype did not affect the preference of the choice with a higher probability of accessing reward (F_(1, 19)_ = 0.007, *P* = 0.936; Fig. 6c). Both control and mutant mice showed consistent preference for the “90%” corner (59.3 ± 0.594% (SEM) and 59.4 ± 0.825% of visits in the compartments offering saccharin, respectively). However, significant effects of time period (F_(2, 38)_ = 5.262, *P =* 0.009) and the interaction of the two factors (F_(2, 38)_ = 3.688, *P =* 0.034) were observed. Interestingly, TIF-IA^DATCreERT2^ mice had a significantly higher preference for saccharin than controls (i.e., the fraction of licks on saccharin bottles, Fig. 6d; F_(1, 19)_ = 4.727, *P* = 0.043), with an average fraction of drinking saccharin at 0.661 ± 0.0436% (SEM) in controls and 0.868 ± 0.0297 in mutants, and an interaction effect on the preference over time (F_(2, 38)_ = 3.290, *P* = 0.048) was observed. Thus, female mutant mice performed a greater number of responses in the task and had higher saccharin preference but were not different in the fraction of choices of the higher probability of reward.

**Figure 6.**
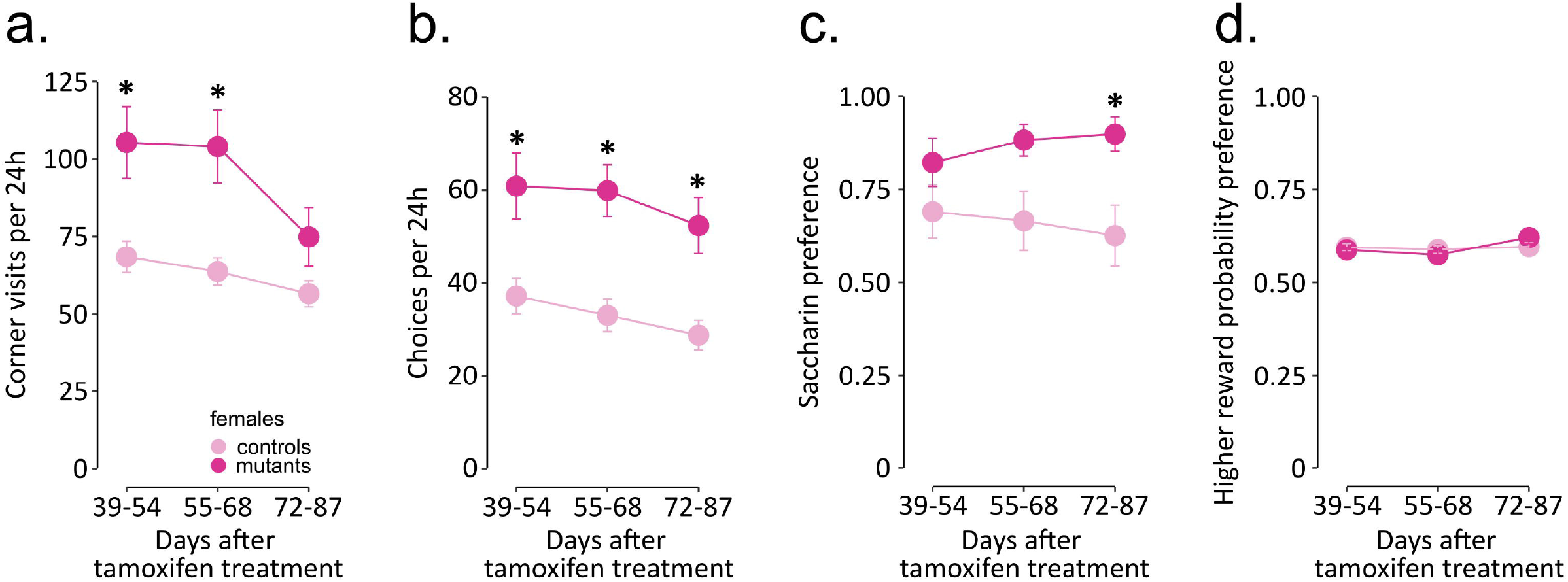
IntelliCage activity in female TIF-IA^DATCreERT2^ and control mice. The graphs show (a, b) activity in the cage, (c) fraction of choices of the higher reward probability and (d) saccharin preference binned over 16-day periods roughly corresponding to the time points from gait testing. Points represent the group mean, and error bars are SEM. Stars indicate significant difference mutant vs. control, *t*-tests with Bonferroni correction for multiple tests, p <0.05.

### Correlations among gait parameters and nonmotor behaviors tested

Correlations between the parameters measured were assessed separately in male and female mice (Fig. 7a & b, respectively, and Supplementary Table S8 with complete correlation matrices). Clear, significant positive correlations existed between motor parameters associated with hind paw print size (area & width; Pearson R: males: R = 0.89, females: R = 0.75) or the hind paw swing speed and cadence (males: R = 0.76, females: R = 0.73), and significant negative correlations existed between cadence and hind paw step cycle (males: R = -0.63, females: R = -0.84) or step sequence AA and coupling of steps between the right paws RF to RH (males: R = -0.51, females: R = -0.55). These pairs of parameters are linked to related gait parameters and, accordingly, were observed very consistently in both male and female mice. In female mice, several significant correlations were observed between motor and nonmotor measures. The base of support for the front paws positively correlated with weight (R = 0.48), which likely reflects the differences in size of the animals (and the increase in size during the experiment). The nonmotor behaviors in females showed strong correlations in closely related measures (e.g., positive correlation between number of visits and number of choices in PRL task, R = 0.87) but also had significant correlations with the gait parameters. The two measures related to the number of corner visits during the probabilistic reversal task had a negative correlation with the base of support for the hind paws (visits: R = -0.41, choices: R = -0.47) and coupling of steps (RH to LF; visits: R = -0.47, choices: R = -0.5). Similarly, saccharin preference was also significantly correlated with the coupling parameter (R = -0.42), as well as with the number of visits (R = 0.51) and number of choices (R = 0.72). In contrast to these observations in females, saccharin preference, weight and the number of active or inactive nose pokes in the instrumental task were not significantly correlated with gait performance in male mice (Supplementary Table S8). Moreover, the number of active and inactive responses performed in the operant sensation seeking task were not closely intertwined (R = 0.32, *P* > 0.05), which implies that the high number of active nose pokes did not depend on the total number of instrumental responses.

**Figure 7.**
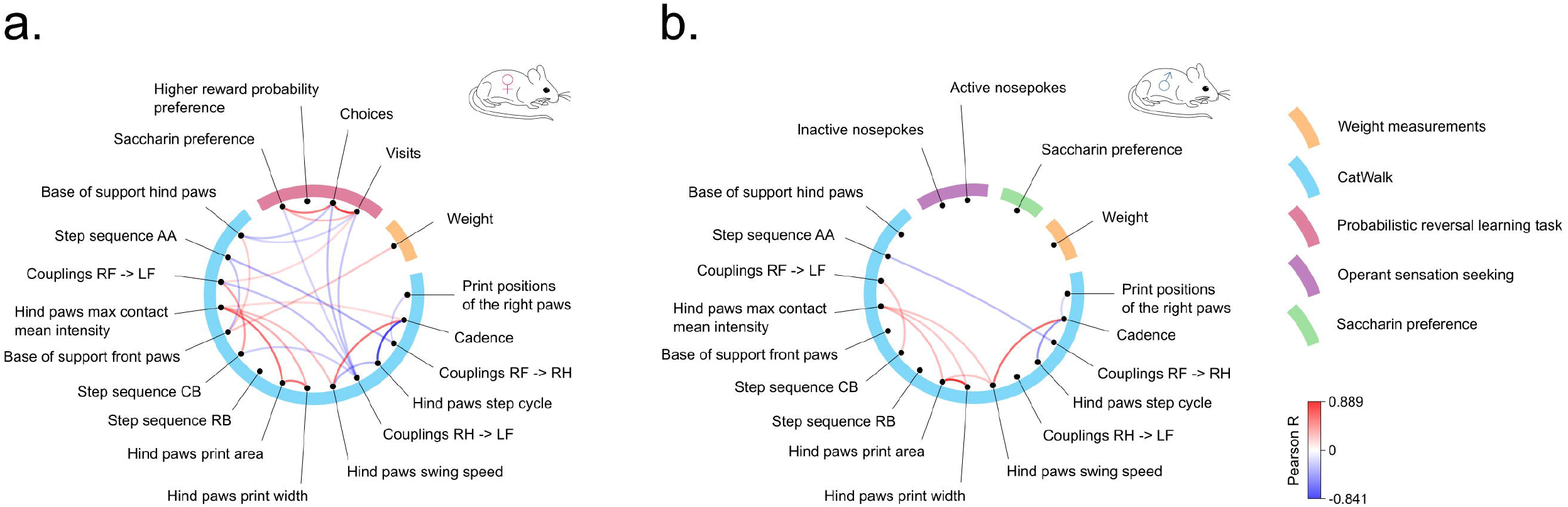
Correlations between gait parameters and nonmotor behaviors. The blue semicircle groups motor parameters, each of them represented by a dot and labeled outside the circle. The remaining colors correspond to the nonmotor parameters. Lines connecting the dots represent significant positive (red) or negative (blue) correlations according to the scale shown.

Taken together, two clear trends emerged in the analysis. First, the results from males and females differed, specifically, the lack of correlation between motor and nonmotor parameters in males. This difference is likely linked to the small number of timepoints for data collection, e.g., 1 observation for 1 block for saccharin preference, which limited the power of the analysis in males and catwalk data in males and females, and the method of testing, with IntelliCage providing a much larger dataset. Second, the distance between hind paws, which was the parameter with the strongest genotype effect, also appeared to have the highest correlation to nonmotor parameter measures, although as noted, significant effects were observed only in females.

## Discussion

We find that the progressive loss of dopaminergic neurons in mice is associated with robustly increased instrumental responding but normal learning of action-outcome contingencies and ability to assess the probability of an action being rewarded. Moreover, this loss was not associated with a decrease in preference for sweet taste. Increased instrumental responding correlated with early changes in gait (the base of stand and step coupling), although these effects were observed only in female mice.

Analysis of gait in TIF-IA^DATCreERT2^ mice revealed only relatively minor alterations occurring up to 12 weeks after inducing the mutation. Previously, the progression of neurodegeneration in the substantia nigra pars compacta was assessed as under 20% at 7 weeks after tamoxifen treatment and 50% at 10 weeks, with loss of dopamine in the striatum reaching 80% at 10 weeks^40^. In humans, the emergence of motor symptoms is reported to occur when the loss of dopaminergic neurons in the substantia nigra exceeds ∼30% and the decrease in dopamine concentration in the caudate is ∼70%^57^. We confirmed that only traces of TH immunoreactivity remained in the striatum of TIF-IA^DATCreERT2^ mice killed at 18 weeks after tamoxifen induction. Considering the robust degree of TH signal decrease in the striatum after 18 weeks of phenotype development and the previously reported impaired motor performance of the mutants at 13 weeks measured on the rotarod apparatus ^39,40^, one could anticipate many differences between mutants and controls in gait at earlier timepoints if detected using a device with a higher sensitivity. Furthermore, worsened performance on the rotarod apparatus could also be a sign of motor dysfunction observable in altered gait parameters^58^. Thus, the observed changes in gait were milder than expected, especially at 11 weeks after mutation induction. Significant effects of the mutation were observed primarily in parameters related to the distance between the hind paws and timing of placing two paws (couplings RH -> LF), although we note that the results were not fully consistent between male and female mice. The results are in partial agreement with those of previous studies that used similar methodology^59–62^. Direct comparison of the parameters is confounded by differences in parameter reduction approaches and, most importantly, the stage of dopaminergic degradation in the animals tested. In a study where transgenic rats overexpressing human α-synuclein were tested for their performance in the Catwalk, the earliest (at the age of 10 to 12 weeks) observed changes in gait were related to hind paw base of stand and hind body speed^60^, which is consistent with our results. Furthermore, the effect of the mutation on weight gain may affect gait and thus represent a confounding factor, although we only observed a correlation between weight and the distance between front paws in females. Moreover, gait impairments in PD patients were found to be only partly remediated by dopaminergic treatment^63–65^, which may also explain the small degree of gait dysfunction observed in TIF-IA^DATCreERT2^ mice. Altogether, we found only relatively minor changes in the gait of TIF-IA^DATCreERT2^ mice, which confirms the limited degeneration of dopaminergic neurons and their impact on gait and excludes movement impairment as a factor that could have affected the testing of nonmotor behaviors.

Loss of olfactory acuity was reported as a frequent prodromal PD symptom^66^. Dopamine signaling plays a major role in the rodent olfactory system^67–69^, and thus, a change in the abundance of dopaminergic neurons could be expected to affect the sense of smell in mice. PD patients have been observed to have an increased number of dopaminergic cells in the olfactory bulb^70,71^, and a direct lesion of dopaminergic neurons in the olfactory bulb prevents the development of hyposmia in 6-hydroxydopamine-treated rats^72^. However, as reported previously, the mutation in TIF-IA^DATCreERT2^ primarily affects midbrain dopaminergic neurons without appreciable effects on the olfactory system^52^. Accordingly, we herein observed no change in TH-positive cell abundance in the olfactory bulb of TIF-IA^DATCreERT2^ mice. Thus, the lack of genotype effects on olfactory acuity in the task requiring a buried treat is consistent with the properties of the Cre transgene in the mutant strain.

We found no decrease in saccharin preference in mutant mice and thus no anhedonia-like behavior that could be interpreted as a symptom of a depression-like phenotype^73^. The significant increase in preference observed in females stems from the larger number of choices performed, and we believe that the volume of sweetened water intake itself is unlikely to represent an increase in reward-seeking, especially since we found no differences between male mutants and controls in this behavior. Previously, we reported that mice lacking NMDA receptors in dopaminergic neurons (the NR1^DATCreERT2^ strain) had a phenotype resembling some depression symptoms; however, these animals also had a normal preference for saccharin^74^. Therefore, the absence of anhedonia alone is not sufficient to rule out other depressive-like symptoms.

The most consistent, nonmotor effect of the mutation was increased activity in instrumental tasks. A higher number of responses was observed in males in the OSS task and in females in the IntelliCage-based probabilistic reversal learning (PRL) task. As noted, this change was not associated with decreased accuracy, and responses on the inactive lever in the OSS and preference for higher reward probability in the PRL task were not affected by genotype. Therefore, TIF-IA^DATCreERT2^ mice were not impaired in instrumental learning. We and others have previously reported that performance in the OSS task is highly sensitive to the loss of metabotropic glutamate receptors (including a selective mutation in neurons expressing dopamine D1 receptors)^44,75^ and to treatment with opioid receptor antagonists^45^. Moreover, the activity of dopamine D2 receptors was previously indicated as a key regulator of sensation seeking^76,77^; thus, we anticipated a change in the behavior of TIF-IA^DATCreERT2^ mice in the OSS. Nevertheless, the observed increase, rather than decrease, is counterintuitive, even more so since PD patients may display lower sensation seeking^78^. The increase in responding could be speculated to reflect compensatory mechanisms that emerge after a partial loss of dopaminergic neurons^79^, and this notion would be consistent with the observed time course – the strongest effect was apparent 7 to 9 weeks after the induction of the mutation, and responding returned to control levels at the 11-week mark.

The lack of genotype effect on worsened performance in the PRL task strongly implies normal executive functions in TIF-IA^DATCreERT2^ up to 11 weeks after inducing the mutation. In humans, PD reportedly causes a shift in sensitivity toward negative outcomes in the PRL, which was influenced by L-DOPA treatment^80,81^, and possibly a general modest impairment in the PRL task^82,83^. However, these observations are subject to continuing discussion, and the higher sensitivity to negative outcomes is especially debated^83^. Due to the strict time constraints and predetermined structure of the PRL tasks applied in humans, an increase in choices like that observed here could not be detected. Moreover, we note that the intervals between choices in the task presented here were usually between 1 and 20 minutes, compared to seconds in typical human PRL tasks. Thus, while the data reported here show no impairment in executive functions in mice during the early stages of dopaminergic neuron degeneration, the increase in instrumental responding certainly warrants investigation in humans, which would necessitate a redesign of the usually applied PRL tasks. This idea is supported by the analysis of correlations between observed motor and nonmotor behaviors. The number of choices and visits performed by female TIF-IA^DATCreERT2^ mice in the PRL task was negatively correlated with hind paw swing speed and base of support. The hind paw swing speed could be considered to have the highest similarity to bradykinesia; thus, the negative correlation to instrumental choices suggests that it may have a common underlying mechanism. Nevertheless, we do note the caveat that the swing speed data have high variability, and instrumental responses in the OSS in males were not significantly correlated with gait parameters.

In summary, we found relatively few changes in behavior in the early stages of dopaminergic neuron degeneration in a genetically modified mouse strain, and the largest effect of genotype was observed in operant behavior. These results do not support a contribution of a partial loss of dopaminergic neurons to impaired executive functions in the prodromal phases of PD; however, they also suggest that increased instrumental responding merits investigation as a potential marker of early stages of disease development.

## Acknowledgements

The study was funded by statutory funds of Maj Institute of Pharmacology Polish Academy of Sciences. AR and JJ were supported by PhD stipends from InterDokMed project no. POWR.03.02.00-00-I013/16 at the time of the study.

## Author contributions

AR, ML, JP, DW, and JRP designed the study; AR performed all behavioral experiments together with JJ, ŁS and ZH; AR performed the histological procedures; AR, ML, and JRP prepared the figures; AR, JJ, ML, JP, DW and JRP analyzed the data; MB and GK generated and provided the genetically modified mice used in the study; AR and JRP wrote the manuscript with help from all the authors.

## Data availability statement

The R scripts used in the statistical analyses are available at the GitHub page of the experiment: https://github.com/annaradli/tif-pd-behavior. Raw behavioral data can be downloaded from the Zenodo webpage of the project: https://doi.org/10.5281/zenodo.7539367.

## Additional information

The authors declare no competing interests.

## Supplementary materials

**Supplementary Figure S1.**
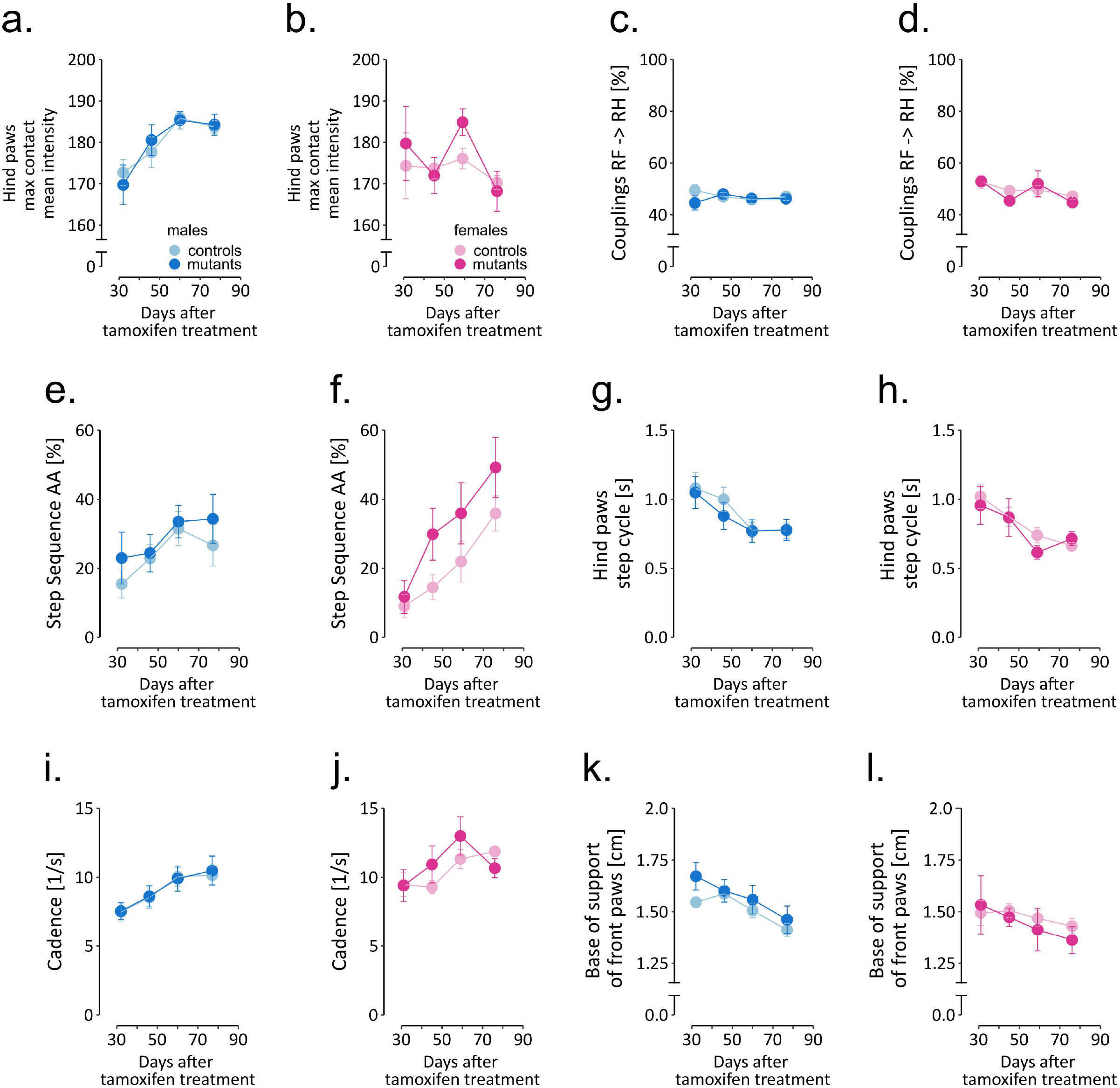
CatWalk parameters implied by the linear regression model that were most closely related to genotype (Figure 4 continued).

Supplementary Table S1. Summary of two-way ANOVAs of all behavioral tests and weight measurements for males and females.

Supplementary Table S2. CatWalk complete raw dataset.

Supplementary Table S3. Correlation coefficients of CatWalk parameters between the left and right paws.

Supplementary Table S4. CatWalk parameters used in linear regression model reduction of data.

Supplementary Table S5. Results of linear regression analysis of CatWalk parameters using the most efficient model.

Supplementary Table S6. Operant sensation-seeking data.

Supplementary Table S7. IntelliCage data summarized in bins.

Supplementary Table S8. Correlation matrices for assessing the relationship between motor and nonmotor functions.

